# Ventralis intermedius nucleus anatomical variability assessment by MRI structural connectivity

**DOI:** 10.1101/2020.08.05.236679

**Authors:** Francisca Ferreira, Harith Akram, John Ashburner, Ludvic Zrinzo, Hui Zhang, Christian Lambert

## Abstract

The ventralis intermedius nucleus (Vim) is centrally placed in the dentato-thalamo-cortical pathway (DTCp) and is a key surgical target in the treatment of severe medically refractory tremor. It is not visible on conventional MRI sequences; consequently, stereotactic targeting currently relies on atlas-based coordinates. This fails to capture individual anatomical variability, which may lead to poor long-term clinical efficacy. Probabilistic tractography, combined with known anatomical connectivity, enables localisation of thalamic nuclei at an individual subject level. There are, however, a number of confounds associated with this technique that may influence results.

Here we focused on an established method, using probabilistic tractography to reconstruct the DTCp, to identify the connectivity-defined Vim (cd-Vim) *in vivo*. Using 100 healthy individuals from the Human Connectome Project, our aim was to quantify cd-Vim variability across this population, measure the discrepancy with atlas-defined Vim (ad-Vim), and assess the influence of potential methodological confounds.

We found no significant effect of any of the confounds. The mean cd-Vim coordinate was located within 1.9 mm (left) and 2.1 mm (right) of the average midpoint and 4.9 mm (left) and 5.4 mm (right) from the ad-Vim coordinates. cd-Vim location was more variable on the right, which reflects hemispheric asymmetries in the probabilistic DTCp reconstructed. The superior cerebellar peduncle was identified as a potential source of artificial variance.

This work demonstrates significant individual anatomical variability of the cd-Vim that atlas-based approaches fail to capture. This variability was not related to any methodological confound tested. Lateralisation of cerebellar functions, such as speech, may contribute to the observed asymmetry. Tractography-based methods seem sensitive to individual anatomical variability that is missed by conventional neurosurgical targeting; These findings may form the basis for translational tools to improve efficacy and reduce side-effects of thalamic surgery for tremor.

**Highlights:**

- Connectivity-based Vim position varied markedly between subjects and from atlas-defined coordinates.
- This positional variability was not related to any methodological confound tested.
- Hemispheric asymmetry was observed in connectivity-based Vim position.
- We hypothesise lateralization of cerebellar functions, such as language, may contribute to asymmetry.
- Knowledge of Vim position variability could help inform neurosurgical planning in the management of tremor.

## 1. Introduction

The ventralis intermedius nucleus (Vim) is a wedge-shaped thalamic nucleus (Figure 1) that is thought to function as a hub for sensory-motor integration (Mai and Forutan, 2012). The Vim nomenclature is part of the Hassler classification and corresponds to the inferior aspect of the ventro-lateral (VLp) nucleus in the Hirai and Jones classification (Hassler et al., 1959; Hirai and Jones, 1989). It contains kinaesthetic neurons, somatotopically arranged along the medial-lateral axis (Asanuma et al., 1983; Strick, 1976; Vitek et al., 1996, 1994), that respond to contralateral muscles and joints (Gallay et al., 2008; Jones, 2007, 1985; Llinás et al., 2005; Sakai, 2013). Sensory information is conveyed to the ipsilateral cerebellum directly via the spinocerebellar tracts (mossy fibres) and indirectly via the inferior olive (climbing fibres). Cerebellar efferents arise from the deep cerebellar nuclei (dentate, interposed and fastigial nuclei); the cerebellothalamic tract (CTT) passes through the superior cerebellar peduncle (SCP) and decussates at midbrain level, forming the brachium conjunctivum. CTT axons pass through and around the anterior contralateral red nucleus without synapsing, terminating in Vim (Gallay et al., 2008). Efferent tracts arise from Vim and project principally to the primary motor cortex (M1) (Jones, 2007), with minor projections to the SMA, pre-SMA and premotor cortex (Sakai, 2013). Collectively, the pathway connecting dentate to motor cortex via the Vim is known as the dentato-thalamo-cortical pathway (DTCp).

**Figure 1.**
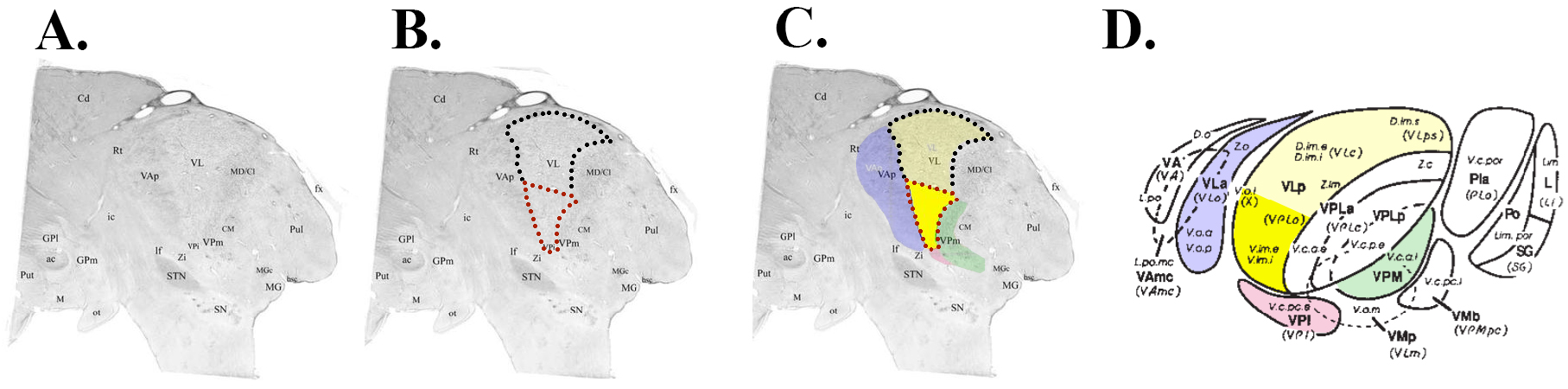
A - Nissl-stained histological sagittal section of a human thalamus at +10.75mm from the midline; B – Outline of VL (dotted line) and position of Vim within VL on this atlas (red dotted line); C. Highlight of surrounding structures (bold yellow = Vim; pale yellow – VL; blue = Ventrolateral anterior; red – Ventroposterior inferior; green - Ventroposterior medial); D - Diagrammatic representation of lateral view of the thalamus showing relationships between thalamic nuclei, colourised as per C. A-C adapted with permission from Ilinsky et al. (2018). D adapted with permission from Gross et al. (2004).

Vim neurons are predisposed to oscillatory firing behaviour that gives rise to tremor (Llinás et al., 2005; Llinas, 1988; Steriade and Llinás, 1988), making this structure a key node in the cerebello-thalamo-cortical “*tremor network*” (Al-Fatly et al., 2019; Muthuraman et al., 2012). In Parkinson’s disease (PD), tremor arises from basal ganglia dysfunction, whereas in essential tremor (ET) it arises from inferior olive and cerebellar nuclei. In both pathologies, the Vim is instrumental in synchronising and propagating abnormal tremor-generating oscillations (Duval et al., 2016; Raethjen and Deuschl, 2012). Consequently, in medically refractory tremor syndromes, the Vim has become a target for stereotactic treatment via deep brain stimulation (DBS) or lesioning techniques (Benabid et al., 1993, 1991; Benabid, 1989; Cury et al., 2017; Hariz et al., 2008; Hirai et al., 1983; Pollak P., Benabid A.L., Gervason C.L., Hoffmann D., Seigneuret E., 1993).

However, the Vim is not visible on conventional MRI sequences and stereotactic targeting in functional neurosurgery typically estimates Vim position from standardised atlas coordinates (Schaltenbrand, 1977; Talairach and J., 1988). These are adapted to individual subjects by using visible anatomical landmarks, such as the mid-anterior and posterior commissural points. Whilst this method provides a reproducible way to identify Vim, it is unable to account for inter-individual variability (Akram et al., 2018); success in stereotactic surgery is dependent on accurate targeting (Horn et al., 2019). To offset this limitation, surgical procedures are often performed with invasive mapping and the patient awake to allow target confirmation via neurophysiological techniques and / or micro-- or macro-electrode stimulation. Such methods are uncomfortable for patients, rely on subjective measures and each surgical trajectory carries a small risk of intracranial haemorrhage leading to neurological deficit or death (Zrinzo et al., 2012). Moreover, intraoperative testing does not always guarantee good long-term tremor suppression as testing can be confounded by implantation oedema. Therefore, there is a clinical need to improve non-invasive pre-operative mapping of Vim to minimise unwanted side-effects, improve outcomes and increase comfort for patients (Chen et al., 2016).

The structural connectivity of Vim and the anatomical properties of DTCp have been proposed as a way to more accurately identify this structure *in vivo* using tractography-based methods (Akram et al., 2018; Coenen et al., 2011; Middlebrooks et al., 2018a; Pouratian et al., 2011; Sammartino et al., 2016; Tian et al., 2018). Tractography reconstructs white matter pathways of the brain using diffusion-weighted imaging (DWI), an MRI modality that measures the diffusive properties of water and has become a widely used tool to investigate the human brain (Mancini et al., 2019). DWI allows the estimation of local fibre orientations at each voxel location; individual white matter pathways can thus be traced by following these orientations from one voxel to the next. A number of methods using structural connectivity to identify Vim apply deterministic tractography but this technique has a number of major limitations (Coenen et al., 2011; Nimsky et al., 2016; Nowacki et al., 2018; Sammartino et al., 2016). It only provides a single tract estimate for each seed, based upon the principal diffusion direction, and is therefore unable to resolve complex fibre orientations such as fibre decussation (Abhinav et al., 2014), it underestimates the size and extent of tracts (Petersen et al., 2017), and generates unreliable results in grey matter regions (Behrens et al., 2003). Given the structural properties of the DTCp, it is clear these issues are highly relevant for Vim targeting. Probabilistic tractography represents an alternative approach as it is able to quantify the uncertainty of multiple fibre-orientations at a voxel-wise level and yields a higher level of precision compared to deterministic methods (Abhinav et al., 2014; Petersen et al., 2017). Several methods using probabilistic tractography to improve localisation and segmentation of thalamic nuclei have emerged (Akram et al., 2018; Behrens et al., 2003; Middlebrooks et al., 2018a; Pouratian et al., 2011; Traynor et al., 2010). While the translational implications for functional neurosurgery are clear, there are a number of confounds that could influence the results, such as head motion (Baum et al., 2018; Le Bihan et al., 2006; Ling et al., 2012; Tijssen et al., 2009) or pre-processing choices used to estimate white-matter connectivity and reconstruct tracts. A pre-requisite for any translational tool designed to aid surgical decision making, is that it should provide an accurate representation of the true underlying anatomy; therefore, possible methodological confounds should be minimised or accounted for in the modelling. Here we follow an established state-of-the-art method to identify Vim *in vivo* using probabilistic tractography. Studying 100 healthy individuals from the Human Connectome Project, we set out to quantify positional variability of Vim and compare tractography-determined coordinates to the surgical standard. We examine the impact of various methodological confounds, including participant motion and regional brain volumes, to establish whether the inter-subject variability reflects true variation in anatomy or simply methodological artefacts.

## 2. Methods and materials

### Ethics statement

This study performed secondary analysis in anonymised human subjects. WU-Minn HCP data was obtained from consenting adults and their families and features that may identify an individual or family unit are safeguarded and not divulged.

### Data and code availability

WU-Minn HCP data is freely available at https://humanconnectome.org. Derived data results and analysis code have been made available in an open repository https://github.com/qmaplab/vim_variability.

### 2.1 Subjects

One hundred healthy, unrelated individuals from the WU-Minn Human Connectome Project were studied (Van Essen et al., 2012). The identification number for each studied individual can be found in our github repository. The minimally processed (Glasser et al., 2013) high-resolution T1-weighted (T1w) and HARDI data were used, including FreeSurfer parcellations for each subject.

#### 2.1.1. Diffusion MRI Imaging Protocol

WU-Minn HCP adopt a multi-shell multi-band spin-echo EPI sequence with three different gradient tables, each table acquired once with right-to-left and left-to-right phase encoding polarities; each gradient table includes 90 diffusion weighting directions and six b=0 acquisitions. Fourier 6/8, field of view (FOV) 210 mm × 180 mm, TR=5520 ms, TE= 89.5 ms. 111 slices were acquired with 1.25 mm thickness and 1.5 mm in-plane resolution, and b-values of 1,000, 2,000 and 3,000 s/mm^2^.

#### 2.1.2 Structural MRI imaging protocol

T1w MPRAGE 3T scans were acquired with TR= 2400 ms, TE= 2.14, TI= 1000 ms and flip angle of 8 degrees, isotropic 0.7 mm voxels and FOV 224 mm × 224 mm. Part of the HCP minimal processing pipeline aligns these images to the diffusion data and re-slices them to 1.25 mm isotropic resolution. These were visually checked prior to processing to ensure good alignment between T1w and diffusion data.

### 2.2 Diffusion data pre-processing

Diffusion data pre-processing was performed using FMRIB Software Library (FSL) v5.0 tools. Diffusion tensor model fitting was done with DTIfit; Bedpostx GPU (Hernández et al., 2013), which applies MCMC to obtain the probability density of the model parameters of the data, was used to estimate three fibers per voxel keeping the remaining FSL parameters at their default settings.

### 2.3 Structural data pre-processing

The structural data pre-processing pipeline used the 1.25 mm isotropic T1w data in diffusion space. SPM12 (https://www.fil.ion.ucl.ac.uk/spm/software/spm12/), running in MATLAB R2017a, was used. *Segment* (Ashburner and Friston, 2005) segmented the T1w images into CSF, grey matter (GM) and white matter (WM). The *Shoot* diffeomorphic registration algorithm (Ashburner and Friston, 2011) used these segmentations to create a group average space with which each individual was aligned. Total intracranial volumes were calculated using the *Tissue Volumes* function. Skull-stripped T1w images were obtained using a mask created from the sum of the GM, WM and CSF segmentations, which was thresholded at 0.1.

### 2.4 Tractography targets

We replicate the method of Akram et al (2018). For this study, the following bilateral regions had already been defined on MNI ICBM 152 non-linear (6th Generation) symmetric standard-space T1-weighted average structural template image (1 mm resolution) (Grabner G., Janke A.L., Budge M.M., Smith D., Pruessner J., 2006): primary motor cortex (M1), thalamus, dentate nucleus. As the DTCp is a decussating fibre pathway, contralateral cerebrum and ipsilateral cerebellum canonical exclusion masks were also used. To accurately estimate warps between the HCP population average space and MNI space, warps between the group average T1w images were estimated using FSL FNIRT, and the results visualised to ensure good alignment. All of the canonical seed, target and exclusion masks were then warped to the group average HCP space. These binary images were then warped, using nearest neighbour interpolation, to the individual subject space using the deformation field generated via *Shoot*.

### 2.5 Probabilistic tractography

FSL’s probabilistic tractography tool for NVIDIA GPUs, Probtrackx GPU (Hernandez-Fernandez et al., 2019), was used. Every voxel was sampled 5,000 times, using the three fiber distributions option, with a curvature threshold of 0.2. The proportion of samples that depart from a seed voxel and reach a target ROI is defined as the probabilistic index of connectivity (PICo). In this study, PICo ranged from 0 to 0.5 and was used to construct a sparse connectivity matrix. To avoid tracking into the ventricles, subarachnoid space and the remainder of CSF spaces, a threshold of 0.2 was used to binarize the CSF segmentation, which was employed as a termination mask. All tractography used the canonical exclusion masks described above. As per Akram et al (2018), the following tractography strategy was used to reconstruct the DTCp:

1. Seed: M1; Waypoints: Contralateral cerebellum and ipsilateral thalamus.
2. Seed: Cerebellum; Waypoints: Contralateral M1 and thalamus.

Both outputs from tractography were then summed to reconstruct the DTCp and used for further analysis. The probabilistic DTCp reconstructions were warped to group average space using the deformation field generated via *Shoot* and trilinear interpolation. These warped DTCps were used to create average DTCps for visualisation purposes (Figure 2).

**Figure 2.**
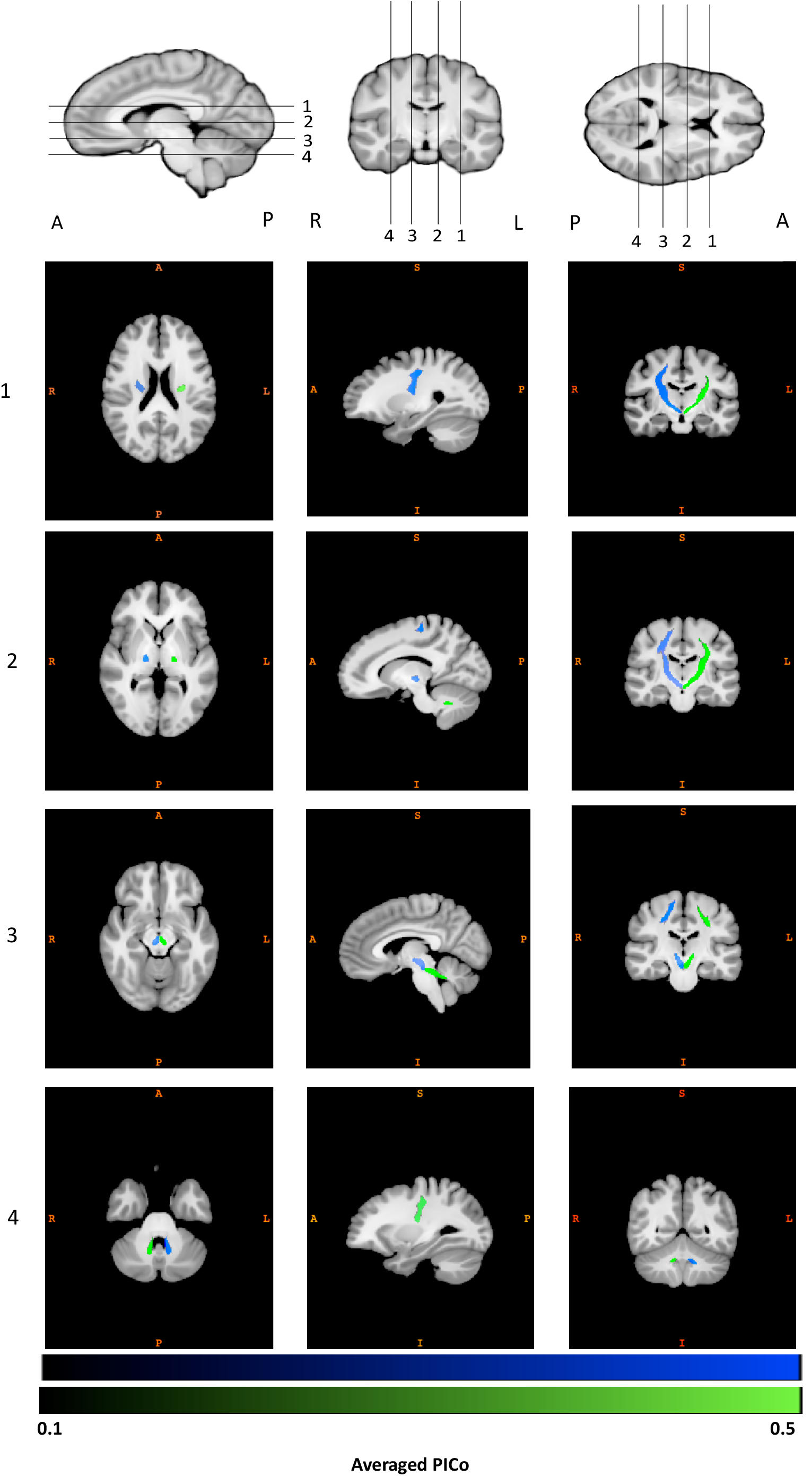
Right (blue) and left (green) group average DTCp windowed between PICo 0.1 – 0.5

### 2.6 Coordinate system

All reported coordinates in this work have been described in the anterior-posterior (AP), medio-lateral (ML) and superior-inferior (SI) directions, following standard stereotactic convention. Voxel-to-world mapping has been used for all reported results.

### 2.7 Tractography-defined Vim localisation

The Vim region was defined as the intersection of the DTCp with the respective thalamic mask for each subject. Probabilistic tractography, despite better resolving grey-white matter transitions and complex fibre architectures, typically yields more false positive tracts; a threshold must be applied to improve specificity, while inevitably reducing sensitivity (Zalesky et al., 2016). While thresholding is a necessity, there is no consensus on the ideal limit and the choice of threshold remains arbitrary (Li et al., 2012). Given the potential variation in connectivity strength with tracking distance and seed volume, both of which may correlate with head size, and that fixed PICo thresholds may bias centroids towards regions with higher anisotropy (i.e. areas likely to have higher PICo values), we opted to use an adaptive threshold for the DTCp-thalamus intersection images by binarizing at 20% maximum connectivity within the intersection region.

The volume of each intersection ROI was calculated in mm^3^. Each ROI was then warped to the HCP group average space, and bwconn in MATLAB used to ascertain the coordinates of each individual centroid in group average space. The average centroid locations were calculated for each side. To compare each individual centroid to the group average coordinates, four displacement metrics were calculated for each ROI: These were the distance (absolute displacement) between the individual and average coordinate in each axis (medio-lateral, anterior-posterior and superior-inferior) and overall Euclidean distance (ED), using voxel-to-world mapping to provide results in mm. To compare differences between average world-locations of centroids between hemispheres, a two-sample repeated measure t-test was used (using the absolute values in the ML axis). To compare differences in variability between sides, a Pitman-Morgan test was used (Gardner, 2001).

Results *P* < 0.05 were deemed to be significant. Centroid locations were summarised using the root mean square of the Euclidean distance, maximum Euclidean distance, and maximum absolute displacement in each axis.

### 2.8 Comparison to the surgical standard

As the Vim is not visible on conventional stereotactic MRI, targeting is generally performed with reference to atlas-defined coordinates. To assess the accuracy of this approach, we calculated the position of the Vim on the HCP group-average template using atlas-reference surgical coordinates. These are defined in relation to the mid-commissural point (MCP) of the anterior commissure-posterior commissure (AC-PC) plane, in relation to the using the following formula (Akram et al., 2018):

- AP direction = (AC-PC length)/3 – 2 mm anterior to PC
- ML direction = Midline ± 12 to 14 mm; (12.5mm used in this study)
- SI direction = 0 mm

We projected this location onto the group average thalamus along with the all tractography calculated centroids using the render toolbox in SPM12. We also calculated the absolute displacement of the tractography centroids from the atlas-defined targets as outlined above and tested for differences along each axis using a one-sample t-test. Moreover, surgical trajectory planning to the atlas-defined Vim was carried out on both sides by an experienced functional neurosurgeon (HA) on HCP group average template using a standard surgical approach and the BrainLab Elements planning platform. In short, entry points were planned on or adjacent to the coronal suture with a lateral angle that allows for trajectories that avoid sulci and the lateral ventricles, maximise the intrathalamic course, and avoid close proximity to the internal capsule (Figure 5).

### 2.9 Investigation of confounds

Methods developed to augment neurosurgical targeting will only be translational if they are robust to artifacts and reflect only true anatomical variability, to ensure safety and efficacy in their application. While HCP data has been pre-processed to correct for motion artefacts, the fact that DWI is extremely sensitive to movement (Baum et al., 2018; Le Bihan et al., 2006; Ling et al., 2012; Muller et al., 2020; Tijssen et al., 2009) drove efforts in this work to establish the influence of subject movement on eventual observed Vim variability.

#### 2.9.1 Methodology confounds

Participant motion was tested using the output from FSL “*eddy*”, contained in the eddy_movement_rms file, which consists of two measures of total movement: the root mean square relative to the first volume acquired, and the immediate preceding volume through the acquisition.

Beyond motion artefacts, the choices of seed and target region can heavily influence the resultant structural connectivity obtained by tractography (Ambrosen et al., 2020). Specifically, tracking target volumes may influence the probability of connectivity, so we tested the influence of the volumes of the M1 and cerebellar masks. The probability of connectivity falls the further from the seed a streamline passes; hence we used total intracranial volumes (TIV) as a surrogate measure of tracking distance across the population.

For each hemisphere, we tested a total of five confound measures of interest (two movement, two ROI volume, TIV) against each of the four displacement metrics (displacement in anterior-posterior, medio-lateral and superior-inferior directions and Euclidean distance). An F test was used to test each confound against the centroid coordinates for each hemisphere.

#### 2.9.2 Superior cerebellar peduncle as an anatomical confound

Based on the known anatomy of the DTCp and the inherent limitations of probabilistic tractography, we identified the superior cerebellar peduncle (SCP) as a potential site that may critically influence our model. This is because the SCP is a narrow white matter structure, 2 to 4 mm in width in healthy adults and less in certain disease populations (Tsuboi et al., 2003), which is surrounded by CSF. In the HCP protocol, it is represented by 2-4 voxels and is the narrowest structure the DTCp fibres must cross. We hypothesised that SCP size may positively correlate with the connectivity metrics in the resulting tracts. For this, we manually segmented the left and right SCP on the group average template using ITK-SNAP. For each individual, we integrated the Jacobian determinant values in these regions to provide a volumetric measure. We then extracted the maximum PICo value for each DTCp and correlated this against the corresponding SCP volume (left DTCp vs right SCP and vice versa).

## 3. Results

### 3.1 Demographics

The cohort consisted of young adults, ranging from 22-33y with an average age of 29.11y (± 3.67y, 54% female) and average brain volume 1.42L (± 0.14L).

### 3.2 DTCp tract reconstruction

Figure 2 shows the group average reconstructed right and left DTCp. Individual probabilistic DTCp reconstructions for five subjects have been provided in the supplementary material.

### 3.3 Tractography-defined Vim

Table 1 summarises tractography-defined left and right Vim coordinates. Overall, the left Vim was located significantly more posterior, inferior and lateral compared to the right. There was more variability in Vim location on the right in the AP and SI axes. Table 2 summarises the dispersion metrics for the tractography defined centroids compared to the group average. Figure 3 shows the principal directions of variability.

**Table 1:**
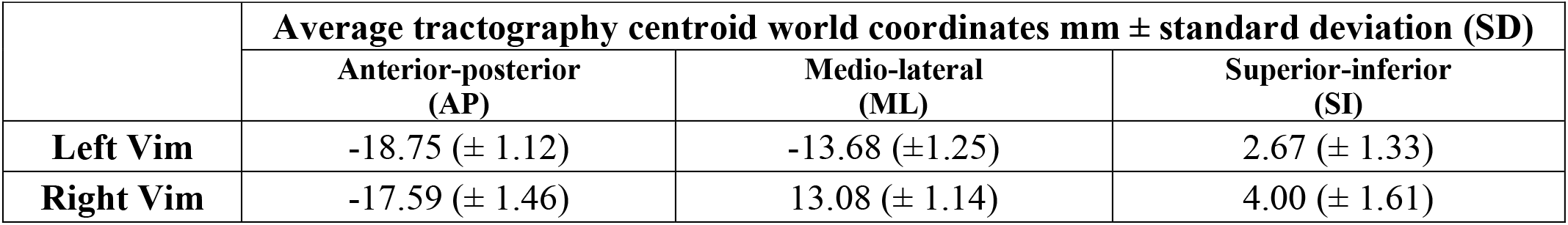
Average (SD) tractography Vim location (world coordinates). Positions posterior to the anterior commissure, left of the mid commissural plane and inferior to axial AC-PC plane are denoted by negative values.

| | Average tractography centroid world coordinates mm ± standard deviation (SD) | | |
| --- | --- | --- | --- |
|  | Anterior-posterior (AP) | Medio-lateral (ML) | Superior-inferior (SI) |
| **Left Vim** | -18.75 (± 1.12) | -13.68 (± 1.25) | 2.67 (± 1.33) |
| **Right Vim** | -17.59 (± 1.46) | 13.08 (± 1.14) | 4.00 (± 1.61) |

**Table 2:**
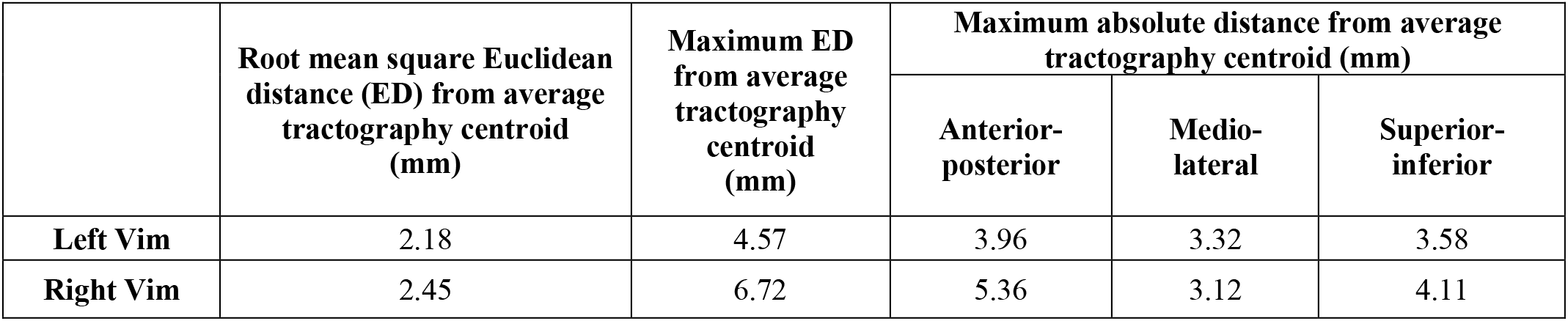
Dispersion metrics for tractography-defined Vim centroids (mm).

**Figure 3.**
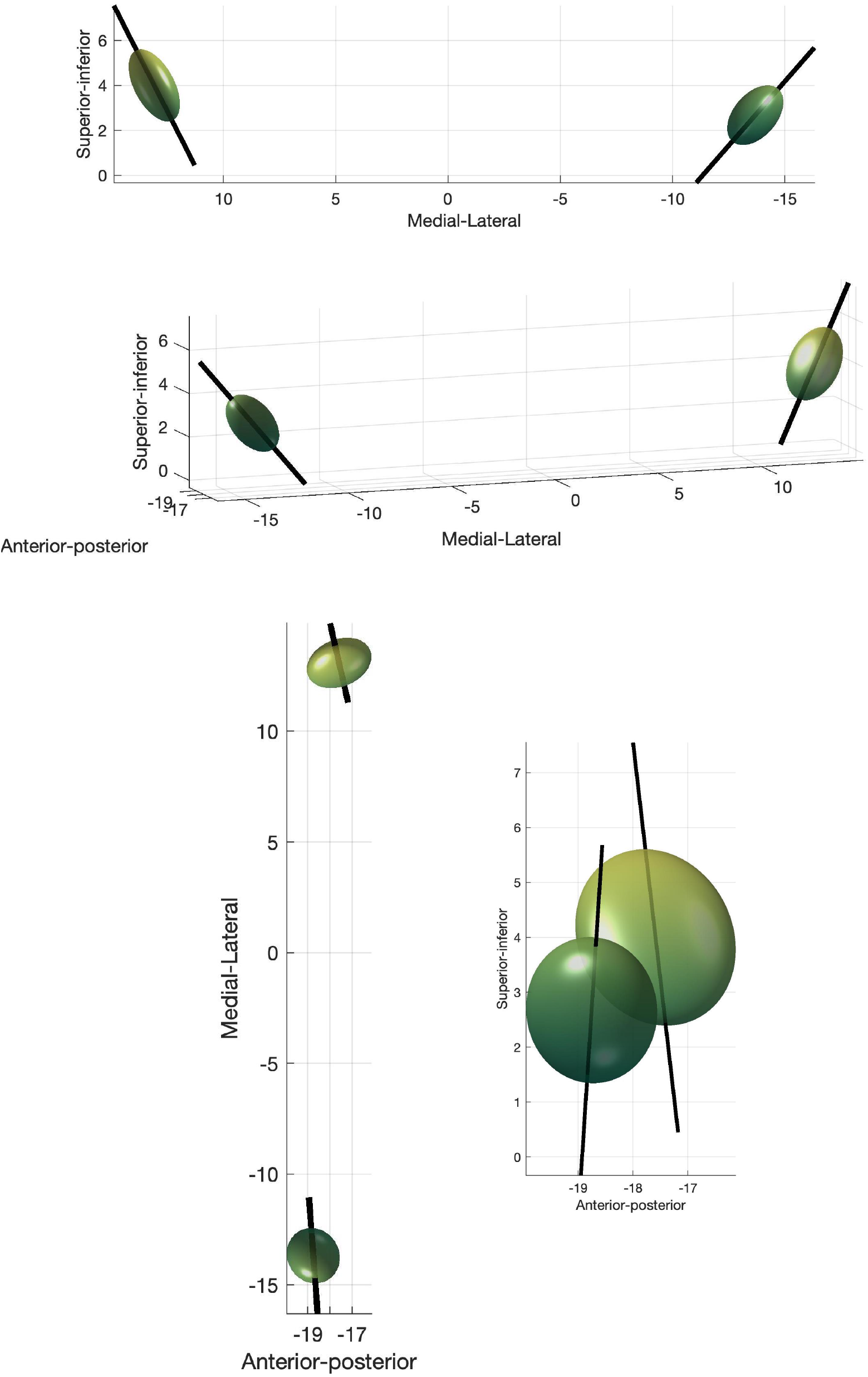
Principal directions of variance for tractography-defined Vim. The eigenvector corresponding to the first principal component is shown as solid line.

### 3.4 Vim coordinates: Atlas-derived versus connectivity-derived

Table 3 summarises the dispersion metrics for the connectivity-defined centroids compared to the atlas-defined surgical coordinates. These were significantly different in all axes, with the surgical coordinates located more medial and inferiorly. The difference was least in the AP plane, located more posteriorly on the right and anteriorly on the left (Figure 4). Figure 5 projects this data onto the thalamus.

**Table 3:** Distances from atlas-defined coordinates.

|  | Root mean square Euclidean distance (ED) from atlas-defined coordinates (mm) | Maximum ED from atlas-defined coordinates (mm) | Maximum absolute distance from atlas-defined coordinates (mm) |  |  |
| --- | --- | --- | --- | --- | --- |
|  |  |  | Anterior-posterior | Medio-lateral | Superior-inferior |
| **Left Vim** | 4.25 | 7.26 | 3.70 | 4.51 | 7.00 |
| **Right Vim** | 5.75 | 9.92 | 6.27 | 4.54 | 8.86 |

**Figure 4:**
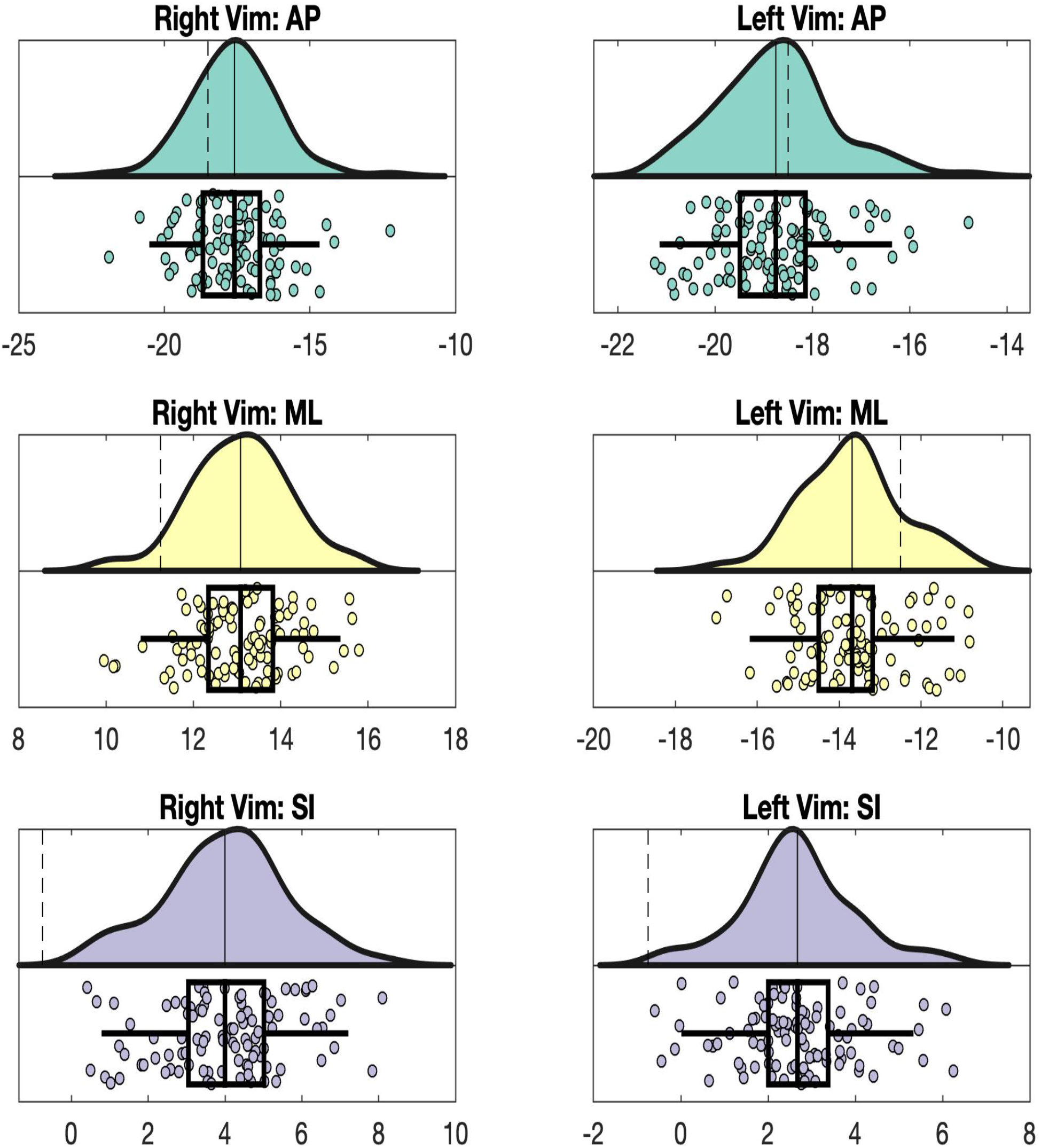
Raincloud plots for tractography defined Vim data. The average tractography (solid line) and atlas (dashed line) defined coordinates are shown on the distributions above. The boxplots below show the original, tractography-defined centroid locations, and mark the average, interquartile range and two standard deviations for each axis (AP, ML and SI), in mm.

**Figure 5:**
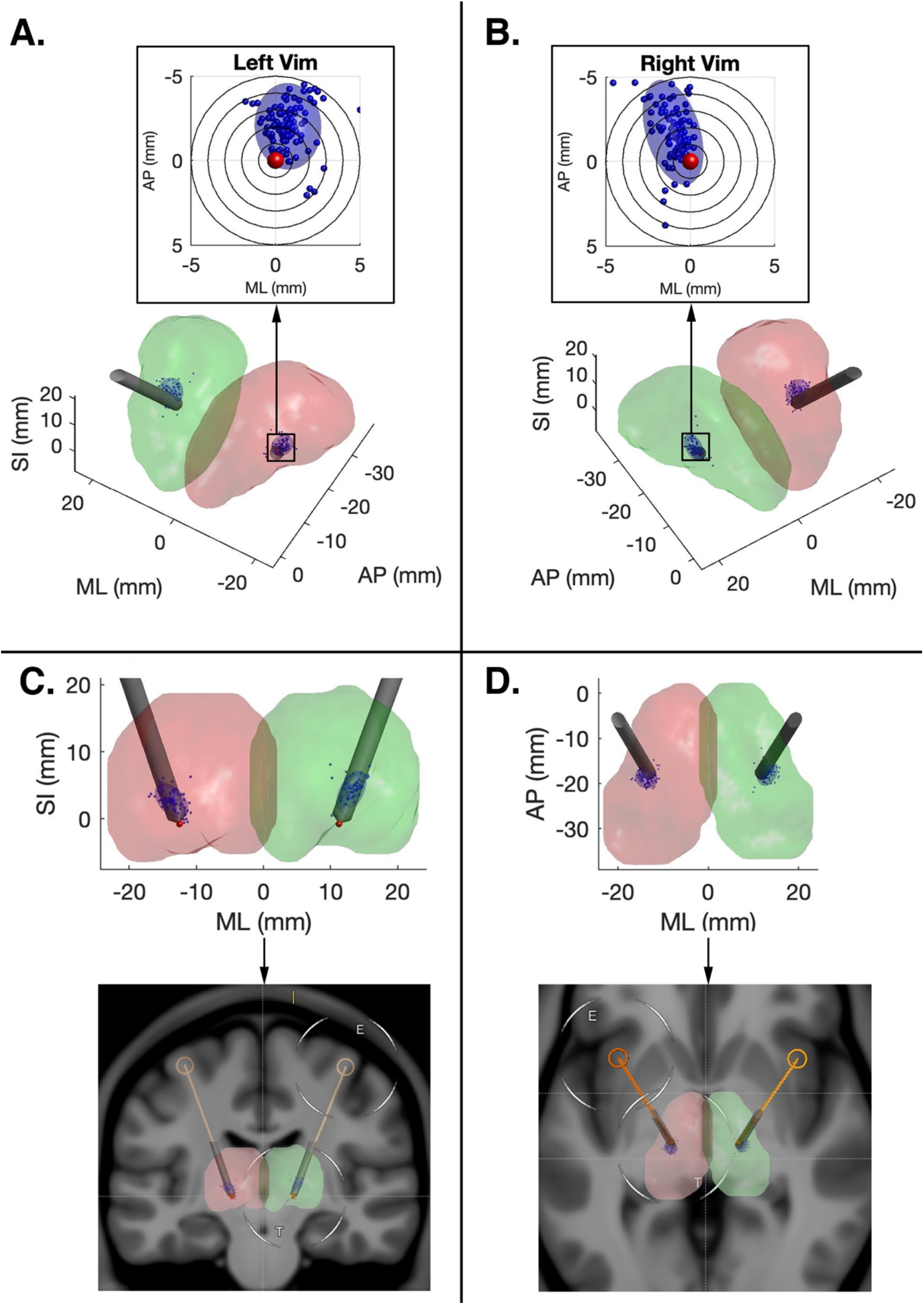
DBS trajectory planning using atlas coordinates. View down bore of left (A) and right (B) thalamic electrode, with individual tract centroids and variance projected orthogonal to the electrode trajectories; Coronal (C) and Axial (D) views of DBS electrode trajectory showing thalamic renderings (top) and original planning MRI (bottom).

**Figure 6.**
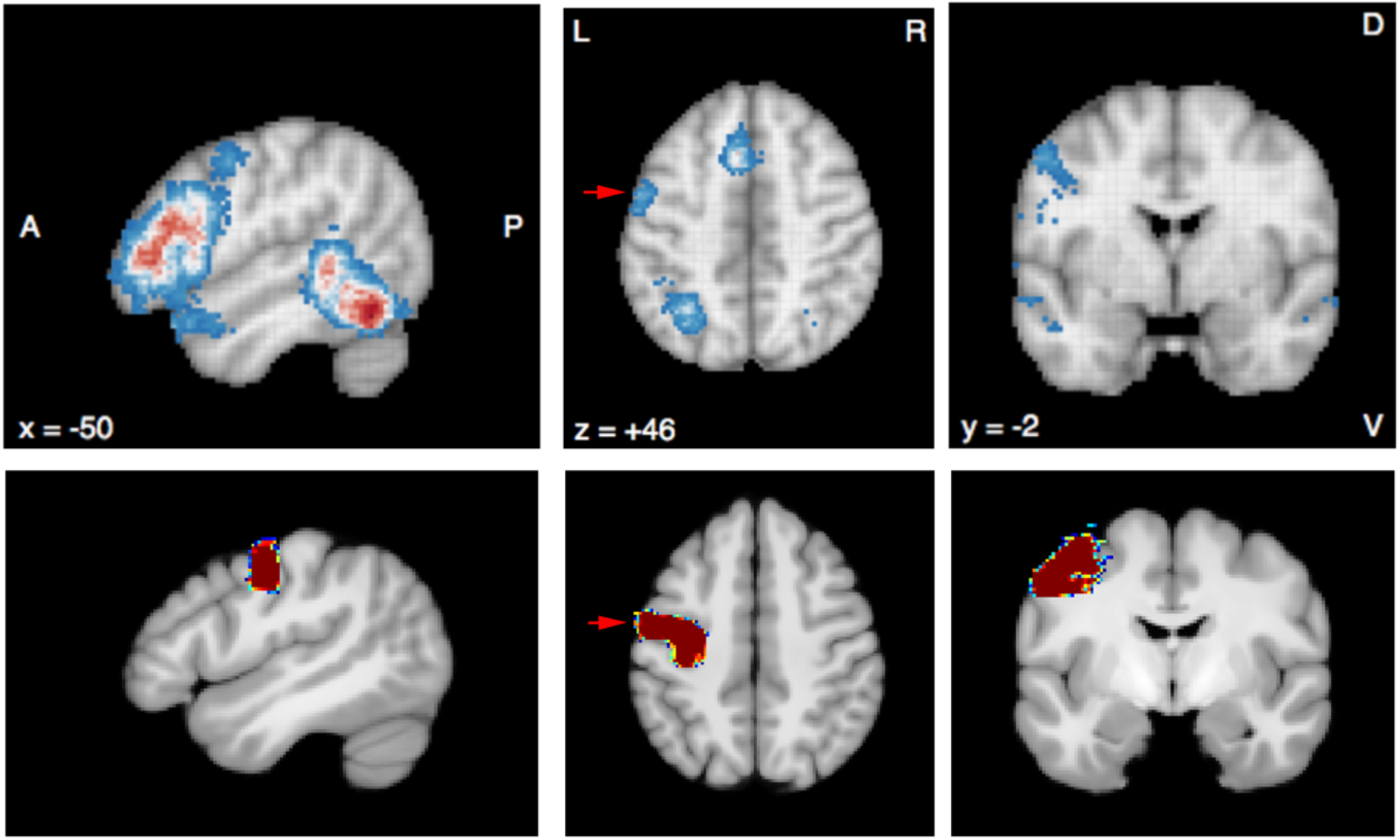
Top row: NeuroSynth fMRI meta-analysis results of 944 studies obtained with search term “word”. Bottom row: left hemisphere M1 mask used for this study (arrow).

### 3.5 Methodological confounds

No significant effects where found for any of the possible methodological confounds (eddy movement parameters, M1 or cerebellar target mask volume, TIV).

### 3.6 Superior cerebellar peduncle

No difference between left and right SCP volumes was found. On the left (i.e. left DTCp vs right SCP), SCP volumes significantly correlate with PICo max, mean and variability. On the right, no correlation exists.

## 4. Discussion

A connectivity-based approach was used to characterize the inter-subject variability of the DTCp in 100 unrelated subjects from the WU-Minn HCP dataset. The DTCp was chosen as it is part of the tremor network that can be targeted in functional neurosurgery in the treatment of severe refractory tremor disorders, such as PD and ET (Anderson et al., 2011). The Vim, not visible on conventional MRI, occupies a central position in the DTCp. The inability to directly visualise the Vim on conventional imaging, and the consequent reliance on atlas-defined coordinates for surgical planning, may lead to suboptimal targeting and poor long-term clinical outcomes (Akram et al., 2018). While demonstrating Vim inter-subject variability, and discrepancy from atlas-defined Vim coordinates, we aimed to exclude a number of possible methodological confounds, including tracking target volume, head size and subject movement in the scanner. None were found to be statistically significant after correction for multiple comparisons. We found that the SCP volume influenced PICo on the left, and therefore may represent a bottleneck for probabilistic tractography; a pontine target, prior to entering this structure, may help generate more consistent results. There were hemispheric asymmetries in the probabilistic DTCp reconstructions, which may be accounted for by intrinsic anatomical idiosyncrasies of lateralized cerebral and cerebellar connectivity.

### 4.1 Vim localisation

*Ex vivo*, it is known that the precise location of Vim varies significantly among subjects (Morel et al., 1997). This represents a significant challenge for the surgical treatment of tremor, where accurately locating Vim in individual subjects is considered to be key for improving therapeutic outcomes (Akram et al., 2018; Pouratian et al., 2011; Tsolaki et al., 2017). Indeed, in Vim DBS it has been shown that an error in lead placement of just 2 mm can lead to a significant impact on efficacy (Papavassiliou et al., 2004). Atlas-derived targeting, the current standard practice for Vim targeting in functional neurosurgery, is widely recognised to be sub-optimal according to both functional and structural connectivity studies (King et al., 2017; Nowinski et al., 2005), confirming the discrepancies between atlas-derived coordinates and individual Vim localization (Akram et al., 2018). In this context, probabilistic tractography represents a promising methodology to demarcate the internal structure of the thalamus at an individual subject level *in vivo* (Akram et al., 2018; Behrens et al., 2003; Middlebrooks et al., 2018b; Pouratian et al., 2011; Traynor et al., 2010).

Exploiting the anatomical properties of the DTCp, it is possible to reconstruct this tract using probabilistic tractography to help identify the Vim *in vivo* (Palesi et al., 2015). Motivated by the translational potential of these methods, we used the method proposed by Akram et al (2018) to better characterise variability of tractography-defined Vim location and determine whether there were any methodological confounds that may influence these results. In the HCP data we found that subject motion, ROI volumes and TIV did not significantly influence results. In line with earlier work (Akram et al., 2018; Middlebrooks et al., 2018b; Papavassiliou et al., 2004; Tsolaki et al., 2017), we demonstrated a significant disparity between tractography-determined and the atlas-determined Vim coordinates. Furthermore, we found that variability in the tractography defined Vim locations was asymmetric, more marked on the right, and that the variation mainly occurred along the anterior-posterior and superior-inferior axes. We also demonstrated that tractography defined Vim locations tended to be located posterio-medially when compared to a standard surgical trajectory. These errors, orthogonal to the standard electrode trajectory (Figure 5), are likely to have the greatest clinical impact. Knowledge of the displacement between atlas and tractography defined Vim may help augment surgical planning in the future, either by using the information to modify surgical trajectories or where this is not possible, for example due to the proximity of eloquent anatomical structures, through the use of directional DBS electrodes.

### 4.2 Cerebellar tract asymmetry

This work found evidence of hemispheric differences in Vim position (both the average position and the positional variability), and also in the relationship between DTCp connectivity metrics and the SCP. Our method to construct Vim centroids is reliant on estimating DTCp via probabilistic tractography, and therefore these results reflect hemispheric asymmetries in the reconstructed tract. Whilst cerebellar functional asymmetry is well established, to our knowledge cd-Vim intrasubject asymmetry has not been previously reported. It is well established that the cerebellum is involved in a wide range of functions, including behaviour, cognition and language (Bostan et al., 2013; Strick et al., 2009). In some of these non-motor activities, marked left-right asymmetries have been demonstrated on fMRI (King et al., 2019) that mirror those seen in the association cerebral cortex, particularly for language generation (Wang et al., 2013). Therefore, one possible explanation for some of the observed DTCp asymmetry may relate to its role in language. Certainly, damage to the right cerebellum has been reported to cause non-fluent aphasias and agrammatic speech. We used the meta-analysis tool NeuroSynth (http://neurosynth.org) to identify language fMRI studies using the search-term “*words*”. This calculated an activation map based on pooled results from 944 studies and revealed a region of significant activation in the inferior portion of the primary motor cortex that fell within the M1 seed mask used in this work (Fig 4). Therefore, some of the asymmetries noted in this work may simply be due to the M1 mask including this speech region.

### 4.3 Limitations

There are a number of potential limitations in this work. There is no gold standard or ground truth for comparison and validation of the tracts generated. However, our objective was to quantify the observed variance and try to exclude possible methodological confounds, as a necessary foundation for developing future translational tools. In the future, we plan to use clinical outcome data from Vim surgery in tremor patients to quantify and refine the accuracy of this approach. Furthermore, it is possible that histological ground truth does not necessarily need to perfectly match when the aim is to generate targeting maps to optimise functional surgical outcomes.

The population used in this work, consisting of young healthy adults, may not accurately represent patients who require surgery for tremor, and it is possible that methodological confounds examined here may have a greater impact in clinical cohorts with movement disorders, necessitating additional steps to control for errors. Furthermore, some measure of ground-truth is required to align any imaging approach with the underlying functional anatomy, as validation. Integrating electrode-specific clinical outcomes from DBS surgery may be one approach to achieve this. This will be the focus of future work, and while the method developed appears to be robust, additional steps may be needed to achieve comparable results in clinical populations; this could include the acquisition of DWI following general anaesthesia to minimise any effect of movement.

Our work used discrete *a priori* anatomical constraints to reconstruct a single well-characterised tract of interest (Akram et al., 2018). Whilst this approach may reduce issues associated with complex-fibre populations and false positive tracts (Maier-Hein et al., 2017), some of the steps, such as choice of target ROIs, masking errors or tract thresholding, may represent additional sources of error influencing the reconstructed anatomy. In probabilistic tractography, thresholding is applied in an attempt to minimise spurious connections, which impact the reconstructed tracts more than false negatives (Zalesky et al., 2016), and the more stringent a threshold the greater the effect in reducing network density (Li et al., 2012). Furthermore, if subjects have different sparsity in their connectivity matrices, thresholding will cause further variations on network metrics, affecting short-term reproducibility of structural connectivity (Tsai, 2018). A potential way to control for this is the application of several connectivity thresholds to characterise the obtained networks (Buchanan et al., 2020). If the sparsity is different within subjects, integration over different connectivity thresholds would represent a way to estimate network metrics relatively reliably (Tsai, 2018). Alternatively, threshold-free approaches have been proposed (Lambert et al., 2017) which avoid this step and may provide more stability.

We found that SCP volume positively correlates with PICo values, an effect that may be accentuated in specific clinical populations. Therefore, choosing more proximal targets, such as the ponto-SCP junction, may eliminate this problem and lead to more reliable results. Finally, there is also a risk that some of the variance could emerge simply through image registration errors, which in this work was primarily achieved using T1w data. Given this, possible approaches to address these issues in the future would be to either adopt non-linear registration models designed for DWI data (Chen et al., 2019; Zhang et al., 2006), or use complementary thalamic segmentation methods to improve the alignment of the internal nuclei (Iglesias et al., 2018; Lambert et al., 2017).

## 5. Conclusions

The Vim is a key target for the surgical treatment of tremor, yet current approaches using atlas-based localisation fail to capture interindividual variability. We have shown that tractography-defined Vim localisation is robust to obvious methodological confounds and can effectively capture anatomical variability *in vivo*. This work provides a foundation for developing non-invasive, translational tools for patient-specific stereotactic targeting in neurosurgery, with the potential to help improve surgical safety, patient comfort and long-term clinical outcomes in these disabling conditions.

## Supporting information

Supplemental digital content

## Abbreviations

AC: Anterior commissure
AP: Anisotropic power
CSF: Cerebrospinal fluid
CTT: Cerebellothalamic tract
CT: Computed tomography
DBS: Deep brain stimulation
DTI: Diffusion tensor imaging
DWI: Diffusion weighted imaging
DTCp: Dentato-thalamo-cortical pathway
ED: Euclidean distance
EPI: Echo planar imaging
ET: Essential tremor
FA: Fractional anisotropy
FOD: Fibre orientation distribution
fMRI: Functional magnetic resonance imaging
FNIRT: FMRIB’s linear image registration tool
FSL: FMRIB’s Software Library
FWE: Family-wise error
FWHM: Full width at half-maximum
GFA: Generalized fractional anisotropy
GM: Grey matter
GPi: Globus pallidus pars interna
GPU: Graphics processing unit
HARDI: High angular resolution diffusion imaging
hiFU: High intensity focused ultrasound
HCP: Human Connectome Project
iMRI: Interventional magnetic resonance imaging
IPG: Implantable pulse generator
L-DOPA: levodopa
MCMC: Markov chain Monte Carlo
MDWI: Mean DWIs with B0 image
MER: Microelectrode recording
MNI: Montreal Neurological Institute
MRI: Magnetic resonance imaging
MS: Multiple sclerosis
PC: Posterior commissure
PD: Parkinson’s disease
PICo: Probabilistic index of connectivity
ROI: Region of interest
SPM: Statistic parametric mapping
SCP: Superior cerebellar peduncle
T1w: T1-weighted
TIV: Total intracranial volume
VBM: Voxel based morphometry
Vim: Ventralis intermedius nucleus
WM: White matter

