## Supplemental digital content for "Ventralis intermedius nucleus anatomical variability assessment by MRI structural connectivity"

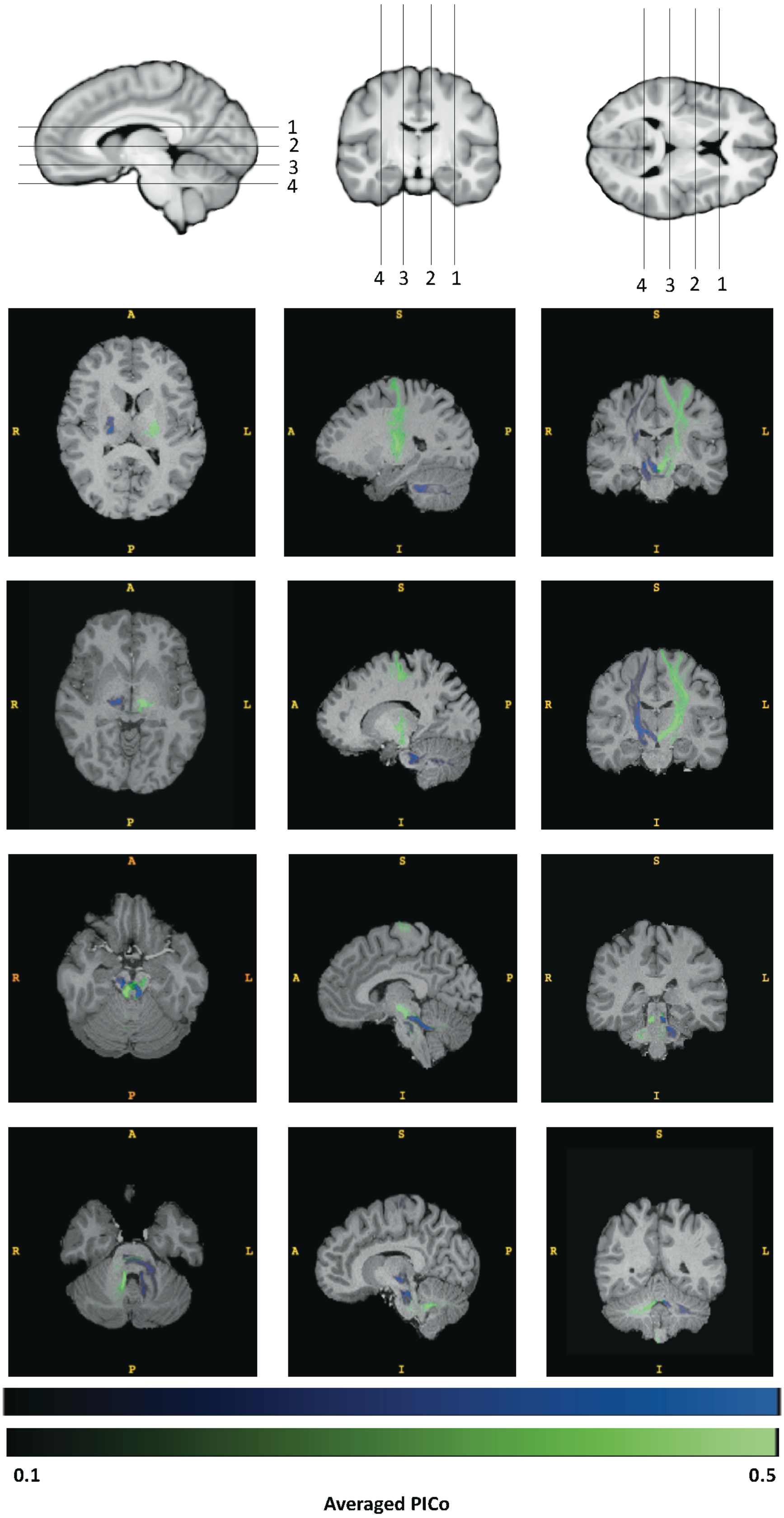


**Figure 7.** Right (blue) and left (green) DTCp for subject 100307, windowed between PICo 0.1 – 0.5

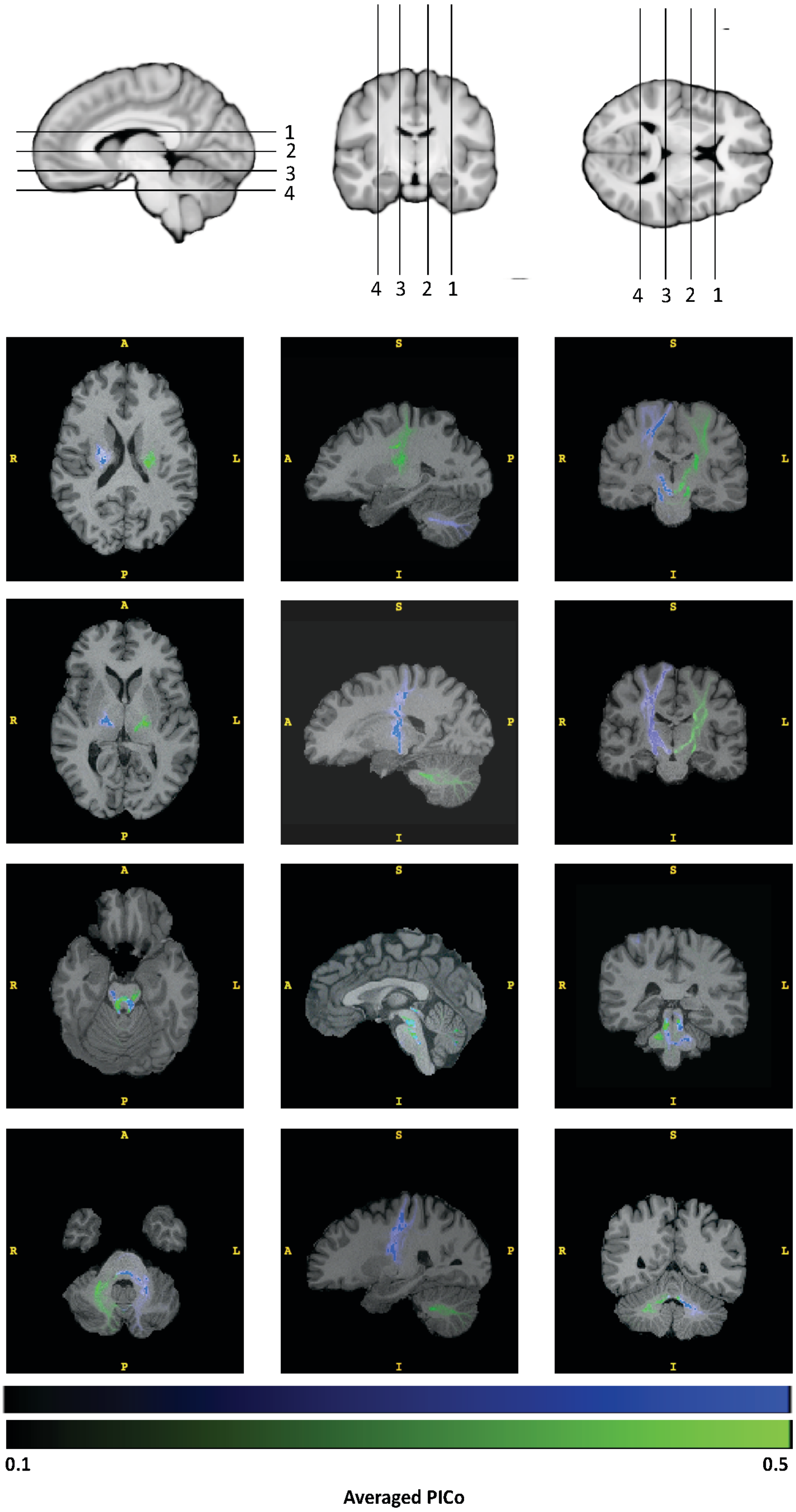


**Figure 8** Right (blue) and left (green) DTCp for subject 100408, windowed between PICo 0.1 – 0.5

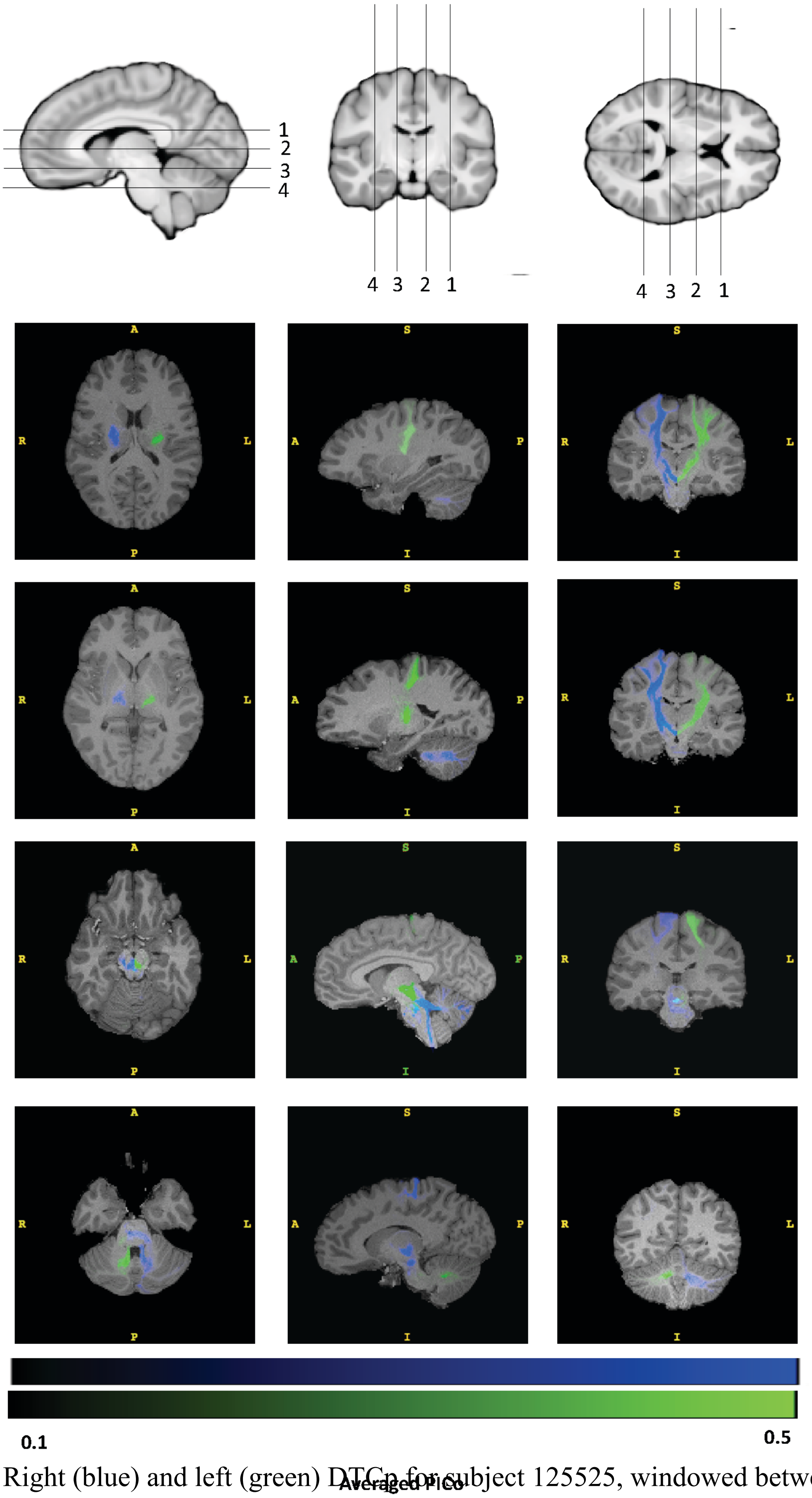


**Figure 9.** Right (blue) and left (green) DTCp for subject 125525, windowed between PICo 0.1 – 0.5

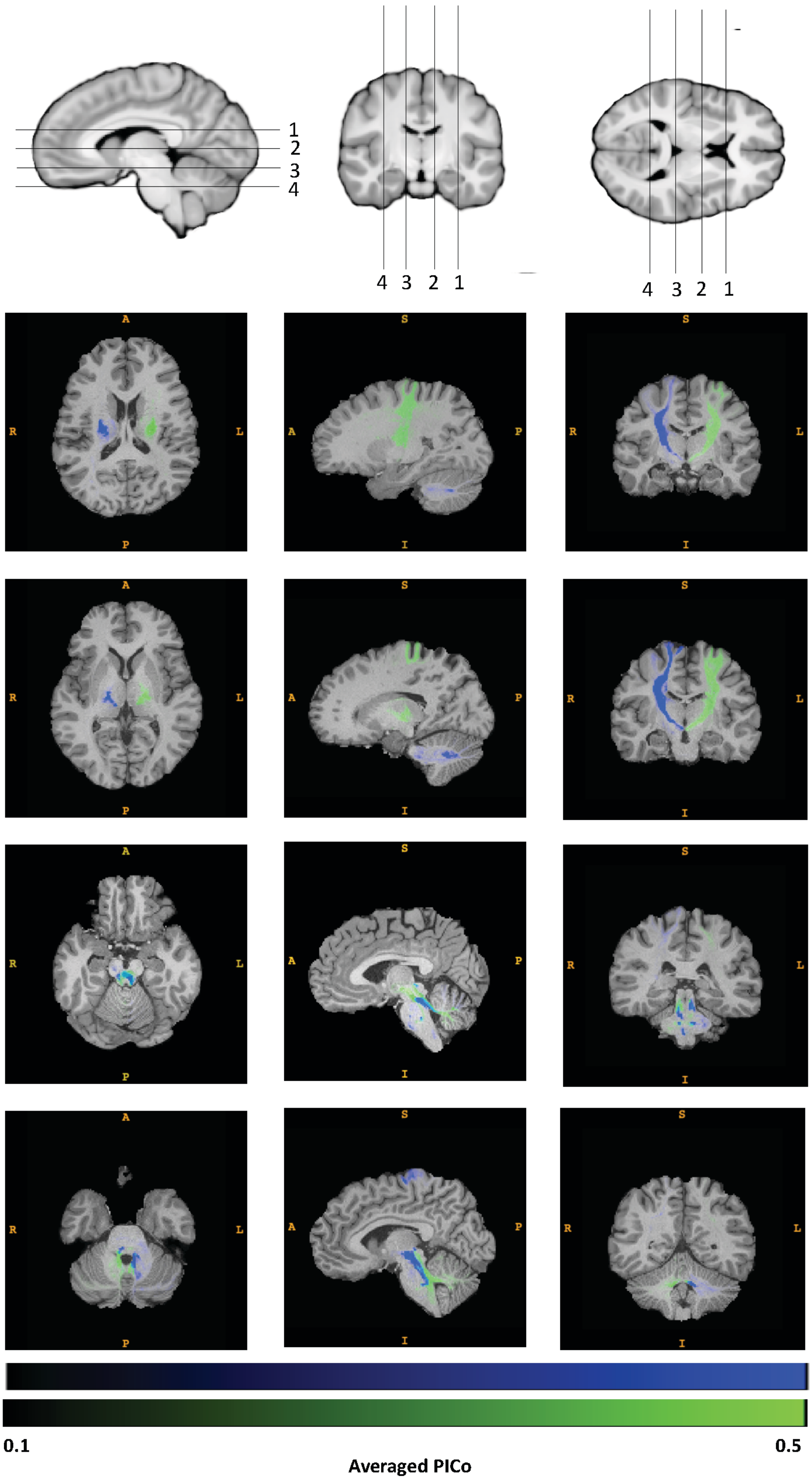


**Figure 10.** Right (blue) and left (green) DTCp for subject 159340, windowed between PICo 0.1 – 0.5

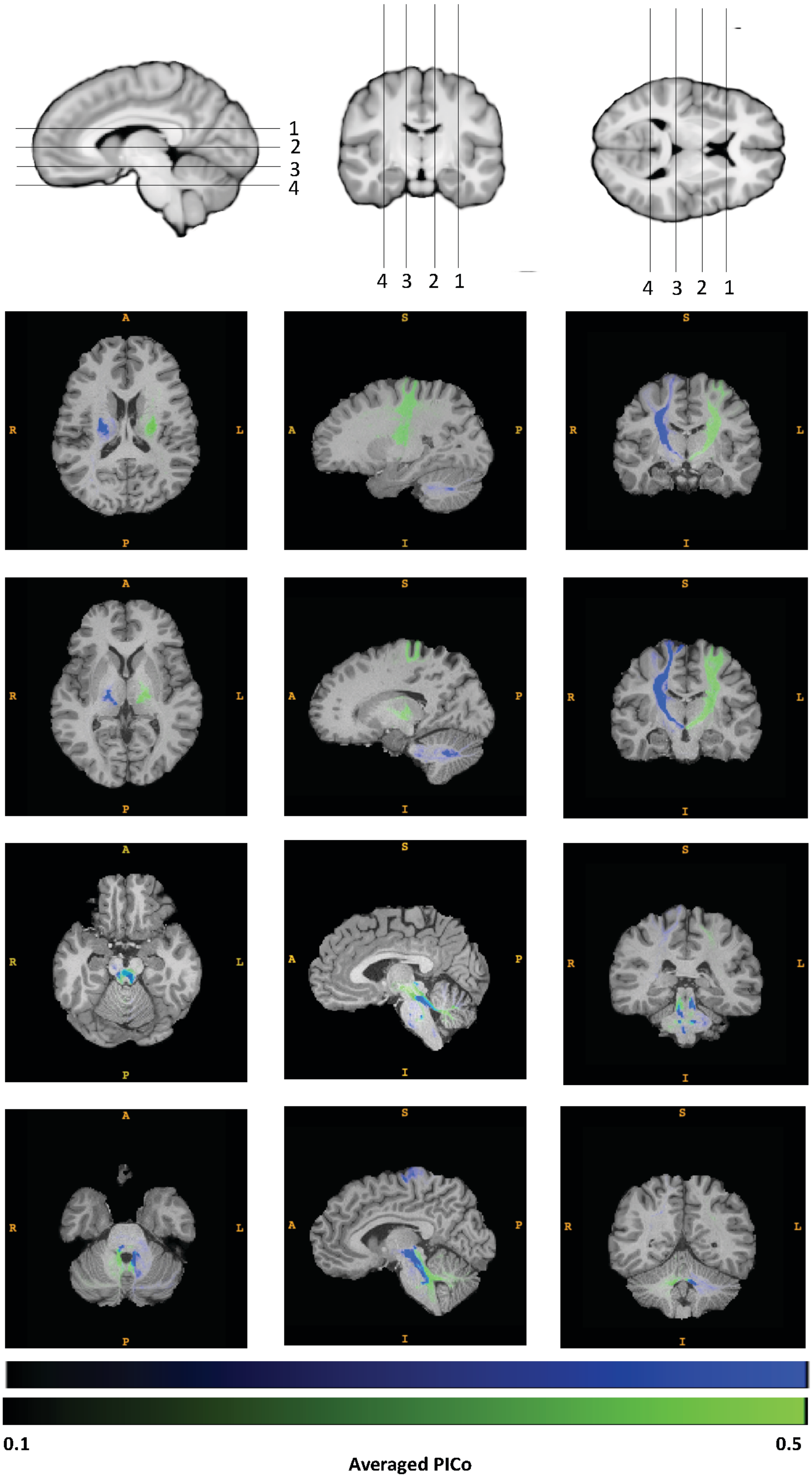


**Figure 11.** Right (blue) and left (green) DTCp for subject 756055, windowed between PICo 0.1 – 0.5
